# Identification of bacteria involved in non-sulfate based hydrogen sulfide production in an aquaculture environment

**DOI:** 10.1101/2024.04.12.589155

**Authors:** Alexandre Nguyen-tiêt, Fernando Puente-Sánchez, Stefan Bertilsson, Sanni L. Aalto

## Abstract

Unwanted microbiological production of hydrogen sulfide (H_2_S) is a major challenge in engineered systems, such as sewage treatment plants, landfills, and aquaculture systems. While sulfur-rich amino acids and other substrates for non-sulfate-based H_2_S production are commonly present, the ability and potential of different microorganisms to perform sulfate-free H_2_S production has not yet been resolved. Here, we reveal the identity, activity, and genomic characteristics of bacteria that degrade cysteine to produce H_2_S in anaerobic enrichment bioreactors seeded with material from aquaculture systems. We compared them with canonical sulfate reducing bacteria and found that both sulfur sources led to microbial H_2_S production, yet with the process being more rapid with cysteine than with sulfate. 16S rRNA amplicon sequencing and metagenomic analysis showed four bacterial families, *Dethiosulfatibacteraceae*, *Fusobacteriaceae*, *Vibrionaceae*, and *Desulfovibrionaceae* to be centrally involved in non-sulfate H_2_S production. Metagenome/metatranscriptome assembled genomes uncovered the main cysteine-degradation pathway mediated by cysteine desulfidase *cyuA* and revealed that some bacteria potentially also use cysteine as a carbon source in sulfate-based H_2_S production.

## Introduction

Hydrogen sulfide (H_2_S) is a volatile sulfur compound present in natural environments (e.g., hydrothermal vents, oxygen minimal zones, marine sediments, hot springs, lakes (1–4); but can also build up in engineered systems (e.g., sewage treatment plants, aquaculture systems, oil and gas processing facilities, landfills (5–7)). While some of this H_2_S is of abiotic origin, microorganisms also play a key role in its production. In both natural and engineered environments, the main group of microbial H_2_S producers is considered to be sulfate reducing microbes (SRM) that use sulfate (SO_4_^2-^) as a terminal electron acceptor for the oxidation of organic or inorganic compounds in order to obtain energy (8). However, H_2_S can also be produced through sulfur disproportionation, where other inorganic sulfur compounds (e.g. thiosulfate, sulfite) are simultaneously oxidized and reduced (9). Microbes can also degrade and desulfurize more complex sulfur (S)-rich compounds, such as cysteine, to produce H_2_S (4,9,10).

Most of the existing knowledge on microbial diversity and enzymes responsible for cysteine degradation comes from the medical sciences (11–13). It has been shown that in microorganisms, cysteine degradation is mainly catalyzed by two different types of enzymes: L-cysteine desulfhydrases and L-cysteine desulfidases, both of which catalyze the decomposition of cysteine into pyruvate, ammonium, and hydrogen sulfide. Enzymes enabling cysteine degradation and H_2_S production can mediate this process as either a primary or a secondary activity. For example, of the four abundant genes coding for cysteine degrading enzymes in the human gut, only *cyuA* (also called *yhaM*) is associated with cysteine desulfidase activity, while the three other enzymes (*mgl*, *malY*, and *cysK*) have other primary functions, but can still use cysteine as a substrate to produce H_2_S as a secondary activity (14–16).

There are some observations on H_2_S production deriving from organic sulfur in nature and wastewater treatment plants. In freshwater lake sediments, H_2_S can for example be produced through degradation of methionine and cysteine under anaerobic conditions (4) and the abundance of proteolytic bacteria capable to break down proteins and degrade the resulting amino acids has been found to correlate strongly with hydrogen sulfide concentration, unlike the abundance of SRM (10). In anaerobic digesters of municipal wastewater treatment plants, the degradation of sulfur-containing amino acids has been found to be more important driver of H_2_S production than sulfate reduction (17), but the identity or metabolic capabilities of the microorganisms involved in this process has not yet been described.

The production of H_2_S is generally in balance with its consumption, as several microbial groups use H_2_S it as electron donor. However, in engineered systems, the prevailing conditions and processes might promote the accumulation of H_2_S if subsequent consumption processes are inhibited. When accumulating, H_2_S rapidly causes serious problems, being flammable, corrosive, and very toxic for eukaryotes even at low micromolar concentrations (18–21) . While the control and prevention of H_2_S-producing bacterial activity has been suggested as a strategy to avoid H_2_S accumulation (22), the current research in engineered environments is mainly focused on the dissimilatory sulfate reduction pathway, neglecting non-sulfate-based H_2_S sources (5,23,24). At the same time, recent studies have found H_2_S production to increase with protein content of sludge collected from a freshwater recirculating aquaculture system (RAS) and from wastewater treatment plant (17,25). High H_2_S production rates also coincide with the presence of the bacteria phylum Fusobacteria (26), which includes the cysteine-degrading, H_2_S producing genus *Fusobacterium* (12,27,28).

In this study, we examined the identity and ecological significance of bacteria responsible for cysteine degradation and H_2_S production in samples collected from a commercial salmon facility, where both sulfate and S-rich amino acids are typically present (marine recirculating aquaculture systems). We performed laboratory reactor experiments and measured the amount of H_2_S produced from cysteine (non-sulfate substrate) and for comparison also from sulfate. Cysteine degradation was further verified by measuring dynamic change in concentrations and coupled stoichiometry of ammonium and volatile fatty acids (VFAs). To shed further light on the organisms involved, we combined 16S rRNA gene amplicon sequencing and in-depth metagenome and metatranscriptome characterization to identify cysteine-degrading H_2_S producers and reconstruct their cysteine-degradation pathways.

## Materials and methods

### Sampling

The starting material for enrichment reactors was collected from a commercial salmon facility in Northern Jutland (Denmark) in May 2022. Three types of starting material were collected; anaerobic activated sludge that represents more degraded organic matter (Aqua-Sample-1), organic material accumulating in the biofilters (biofilter backwash) that represents fresh organic matter (Aqua-Sample-2), and suspended carriers with biofilm from the biofilters (Aqua-Sample-3). For each starting material, three samples of 2 ml were collected and stored at -20°C for further analysis of initial microbial community composition.

### Enrichment of potential H_2_S-producers

To examine the different routes for H_2_S production, two different enrichment conditions were used: media with cysteine to enrich cysteine-degrading H_2_S producers and media with sulfate to enrich traditional *dsrA/B*-carrying H_2_S producers (Supplementary Table S1). For each enrichment condition and type of starting material, three 2-liter anaerobic batch reactors were filled with 100 ml of starting material (corresponding to total chemical oxygen demand of appr. 0.5 g O_2_/l) and 900 ml of corresponding media. Reactors were sealed with screw caps fitted with two ports for sampling purposes and were subsequently flushed with N_2_ for 15 min to create anaerobic condition. The enrichment reactors were operated for 4 weeks at 15°C with continuous magnetic stirring at 150 rpm. Every week, the magnetic stirring was stopped for 15 minutes to let enriched biomass settle, after which 800 ml of media was removed, and the reactor was filled with 800 ml of fresh media. At 1, 5, 7, 10, 14, 21, and 28 days of incubation, 50 ml of biomass sample was collected from stirred reactors for analysis of total dissolved sulfide (TDS) and sulfate (only from sulfate treatment reactors). Samples for microbial community/metagenomic analysis were collected from all reactors on days 0, 14, 21, and 28, and subsequently stored frozen at –20°C. Cysteine reactors were kept running until day 76 for a final sampling occasion.

### H_2_S production rates and pathways of cysteine-enriched microbial communities

To confirm the production of H_2_S via cysteine degradation, we conducted a series of 48-hour experiments using biomass collected from the cysteine-enriched reactors. For the first two experiments, enriched biomass was harvested from Aqua-sample-1 and 2 cysteine-enriched reactors at days 28, 35, and 76 by gentle centrifugation (4,000 rpm for 15 min at 4°C), resuspended in cysteine media (“cysteine-fed”; n=3) (10% v/v) and transferred into 1-liter glass bottles. Incubation bottles were sealed with screw caps fitted with two ports for sampling purposes and were subsequently flushed with N_2_ for 15 min to create anaerobic condition. The incubations were maintained for 48 hours at 15°C with stirring at 150 rpm. A biomass sample of 5 ml was collected in the beginning (0 h) and after 24 and 48 hours for analysis of TDS production and after 24 hours for RNA sequencing (only for experiment 1). In experiment 3, cysteine-enriched biomass from Aqua-sample-2 was collected in the same way, but the biomass was resuspended in cysteine (“cysteine-fed”; n=3) or sulfate media (“sulfate-fed”; n=3) and divided into three 250 ml bottles to create a replicated setup. A biomass sample was collected in the beginning (0h) and after 24 and 48 hours for measuring concentrations of TDS, total ammonia nitrogen (TAN), VFAs (formate, acetate, butyrate, propionate, and valerate), and sulfate (only sulfate-fed bottles) and after 48 hours for RNA. In experiment 4, Aqua-sample-2 biomass was used and resuspended to either cysteine media or media with cysteine and sulfate (Supplementary Table S1). TDS and RNA samples were collected for experiment 3.

### Chemical analysis

Immediately after each sampling, pH, and temperature were measured from the collected biomass sample using a portable meter (Hach HQ40d, Hach Lange, Germany). For measuring total sulfide concentration, aliquots of 0.1 to 2 ml were fixed with ZnAc (10% v/v) and analyzed as in (26) using a spectrophotometer (DR 3900, Hach Lange). The concentration of H_2_S was calculated from TDS using temperature, salinity, and pH according to (29). For measuring VFAs, sulfate, and TAN concentrations, biomass samples were first centrifuged at 4,500 rpm for 15 min at 4°C. Supernatants were then retrieved and filtered through 0.2 µm syringe filters (Filtropur S, SARSTEDT, Germany), and kept at +4°C until analysis. Samples for VFA determination were further fixed by adding 1% v/v of 4 M sulfuric acid (H_2_SO_4_, Merck Millipore, Germany). The concentrations of VFAs and sulfate were determined using a 930 Compact IC Flex 1 with a Metrosep A supp 7 -250/4.0 column coupled with an 887 Professional UV/VIS detector (Metrohm, Sweden), using 0.1 M H_2_SO_4_ as suppressor and 3.6 mM Na_2_CO_3_ as eluent and TAN concentration was determined by 930 Compact IC Flex system (Metrohm, Switzerland) with 887 Professional UV/VIS Detector (Metrohm, Switzerland).

### DNA and RNA extractions

DNA was extracted using the DNeasy PowerLyzer™ PowerSoil DNA Isolation Kit (Qiagen, Germany) according to the manufacturer’s instructions and DNA concentration in extracts was measured with Qubit™ dsDNA High Sensitivity (HS) assay kits and Qubit 2.0 Fluorometer (Thermo Fisher Scientific, Waltham Massachusetts, US). RNA was extracted using the RNeasy kit (Qiagen, Germany) according to the manufacturer’s instructions and the concentration was measured with Qubit™ RNA High Sensitivity (HS) assay kits and Qubit 2.0 Fluorometer.

### Microbial community composition in the enrichment reactors

Microbial community composition was studied by 16S rRNA amplicon sequencing of Aqua-Sample-1 and 2 at day 14, 21, and 28 (sulfate-enriched condition) and 14, 21, 28, and 76 (cysteine-enriched condition) as described in (30). For comparison we also included the starting material used for these incubations in the analysis. Aqua-Sample-3 was discarded from further DNA/RNA-based analysis due to difficulties obtaining representative samples for the biofilm biomass. The detailed step by step protocol for the two-steps PCR procedure can be found in the protocols.io repository (dx.doi.org/10.17504/protocols.io.468gzhw). Hypervariable regions V3-V4 of the 16S rRNA gene were amplified with primers 341F (5′-CCTACGGGNGGCWGCAG-3′) and 805F (5′-GACTACNVGGGTATCTAATCC-3′), resulting in an average amplicons size of 460 bp. The amplicon libraries were sequenced with Illumina MiSeq v3 sequencing chemistry and 2X300 cycles. In total 700,805 reads were obtained from the 18 samples. Raw sequencing reads were processed with QIIME 2 (Release 2022.8/ Version 2022.8.3; (31)). Raw sequences were demultiplexed using the q2-demux plugin and DADA2 (via q2-dada2) was used for trimming, denoising and dereplicating to produce amplicon sequence variants (ASV) (32). Taxonomy was assigned to ASVs using a Bayesian Naïve classifier pre-trained against the Silva database (v138.1; (33)). Files generated in QIIME 2 were transferred to R (version 4.2.1; (34)) for statistical analysis and visualization using packages “phyloseq” (35) and “ggplot2” (36) after singletons and unassigned reads were removed.

### Metagenome sequencing, binning and metatranscriptomic analyses

Shotgun metagenome sequencing was carried out for four samples from day 28 (Aqua-Sample-1 and Aqua-Sample-2 from cysteine and from sulfate-enriched conditions) and two samples collected on day 76 (only cysteine-enriched conditions). Metagenomic shotgun libraries were prepared using the MagicPrep system (Tecan Genomics) according to the manufacturer’s instructions. Short read Illumina NovaSeq 6000 sequencing on an SP flow cell was performed at the SciLifeLab National Genomics Infrastructure in Uppsala (Sweden). Assembly, functional and taxonomic assignment of genes, and bins were obtained by using the SqueezeMeta automatic pipeline (version 1.6.0 (37)). In brief, all metagenomes were pooled and co-assembled with MEGAHIT (38) in order to obtain a single reference assembly. Functional and taxonomic annotation was obtained using the DIAMOND software (39) with three different databases: (i) GenBank nr for taxonomic assignment, (ii) eggNOG v4.5 for COG/NOG annotation (40), and (iii) the KEGG database for KEGG Orthology (KO) annotation (41). A final gene classification was carried out using the PFAM HMM database and the HMMER3 software (42). Coverage and abundance of genes and contigs were estimated with Bowtie2 (43), where custom scripts from the SqueezeMeta pipeline subsequently computed and normalized the results. Binning was performed with CONCOCT (44) and MetaBAT2 (45), and the resulting MAGs were combined into a single non-redundant set with DAS Tool (46). Bin refinement was performed outside SqueezeMeta using RefineM (47) to identify and remove outlier contigs with abnormal tetranucleotide signals, GC patterns and coverage. Contigs were also classified from their taxonomic assignments against the latest reference database provided by RefineM (gtdb_r95_protein_db.2020-07-30.faa) and if the affiliation was different from the bin, the contigs were removed. GTDB-tk v2.3.0, data release R214 (48), was used with default settings to assign taxonomy to each MAG. CheckM (49) was used to provide information about completeness, contamination, and strain heterogeneity. MAGs surpassing a 70% completeness threshold with less than 10% contamination were kept for further analysis. For metatranscriptomic analysis, RNA was extracted from the samples collected in the H_2_S-production experiments 1, 3, and 4. Metatranscriptomic libraries were prepared using the Illumina Stranded Total RNA library preparation kit with ribosomal depletion using Ribo-Zero Plus (Illumina Inc.). Short read Illumina NovaSeq 6000 sequencing on an SP flow cell of 2X150 bp was performed at the SNP/SEQ facility of the National Genomics Infrastructure in Uppsala. Only metagenomics reads were used for assembly and binning, while both the metagenomic and metatranscriptomic reads were mapped back to the assembly for abundance estimation. This was achieved by adding the flags “noassembly” and “nobinning” to the metatranscriptomic libraries in samples file used as an input for SqueezeMeta. Metagenome and metatranscriptome data were further analyzed and visualized using R (version 4.2.1; (34)) and package “SQMtools” (50). The number of transcripts per million (TPM) has been recalculated based on the total number of reads assigned to the BIN rather than by sample before creating gene expression heatmaps. Differences in gene expression between cysteine and sulfate condition were analyzed with DESeq2 (51). DeSeq2 was chosen as it has been shown to provide accurate results even when the number of replicates is low (down to two replicates per condition as long as there are three conditions or less, as in this study (52, 53). Following the recommendations in (52) for using DESeq2 with three or less replicates per condition, a log2 Fold Change threshold of 0.5 was applied in addition to the standard 0.05 adjusted p value significance threshold.

## Results

### H_2_S production in the enrichment reactors and incubation experiments

During the 4-weeks enrichment period, reactors enriched with cysteine started producing H_2_S earlier and faster as compared to sulfate-enriched reactors: H_2_S concentrations of 51 - 82 mg H_2_S/l were reached after 5 days, while sulfate-enriched reactors reached this concentration only after 14 - 28 days (Fig. 1). In cysteine-enriched Aqua-Sample-2 reactor, H_2_S concentration decreased between days 14 and 21, most likely due to oxygen contamination, which led to the consumption of the accumulated H_2_S by sulfide-oxidizing bacteria. H_2_S production had recovered by day 28 (Fig. 1).

**Figure 1.**
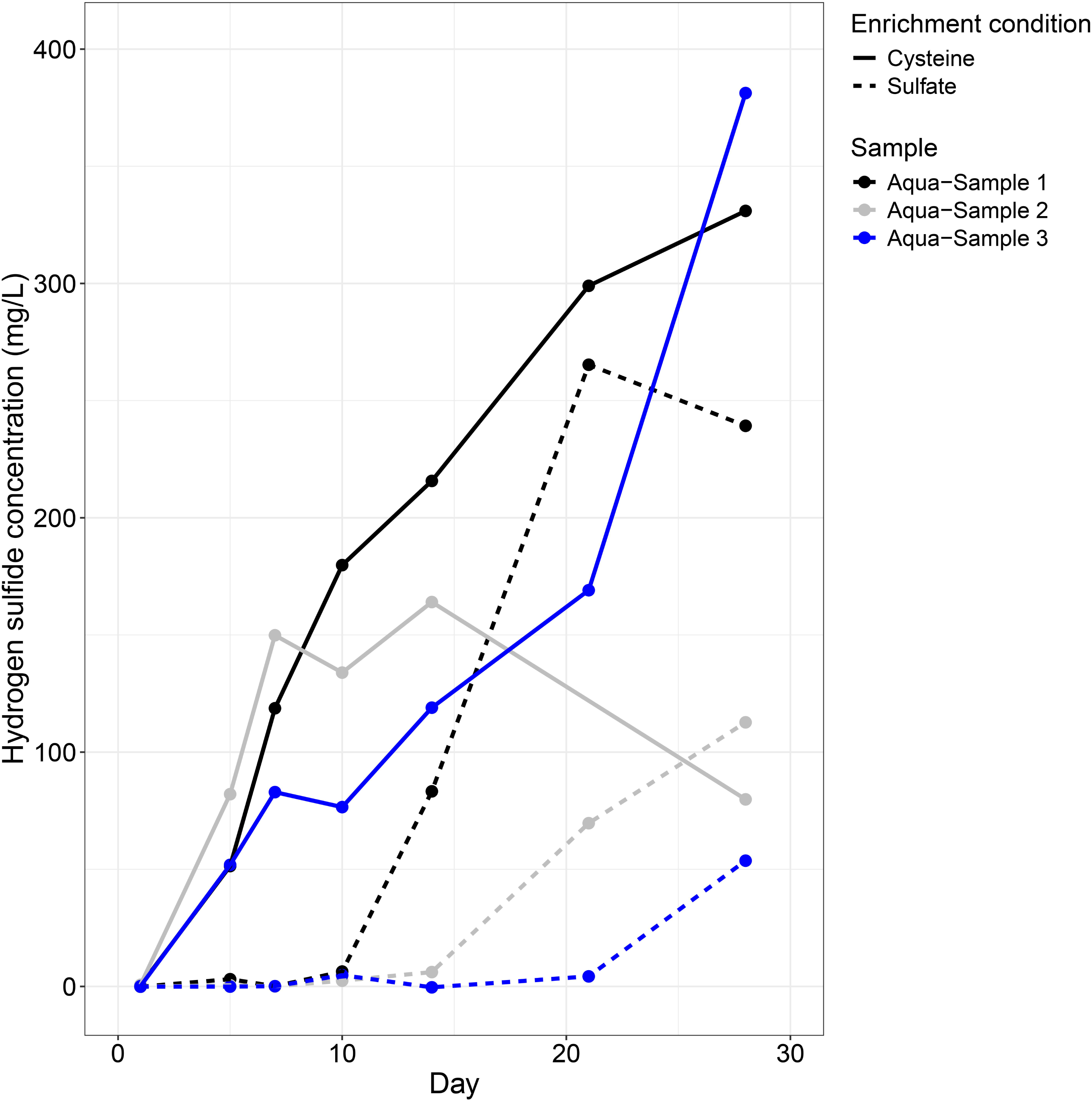
Hydrogen sulfide concentrations (mg/l) under two different enrichment conditions (solid line = cysteine-enriched, dotted line = sulfate-enriched) from three samples, Aqua-Sample-1 in black, Aqua-Sample-2 in grey and Aqua-Sample-3 in blue.

In the 48-hour experiments with cysteine-enriched biomass fed with either cysteine or sulfate media, we observed an increase in H_2_S concentrations and production rate of 9.99 ± 7.86 mg H_2_S/l/d across Aqua-Sample-1 and Aqua-Sample-2 material when using cysteine media (Table 1). At the same time, both TAN and acetate concentrations increased, showing that both compounds were being produced in the cysteine-fed incubations. The formate concentration decreased fast in both cysteine and sulfate-fed incubations, while there were no significant changes in the concentrations of other VFAs (Supplementary Table S2). For sulfate-fed incubations, no production of H_2_S, ammonium, or acetate (Table 1), or consumption of sulfate (data not shown) was observed. The one sample fed with a combination of cysteine and sulfate had a significant increase in H_2_S and consequently a high H_2_S production rate of 23.9 mg H_2_S/l/d.

### Microbial community composition in the enrichment reactors

In the original material collected from recirculating aquaculture system, a large proportion of the microbial community was comprised of families outside the 25 most abundant ones (Fig. 2). Of the 25 most abundant families, *Flavobacteriaceae*, *Candidatus Campbellbacteria*, *Rhodobacteraceae*, and *Rubritaleaceae* were detected in the original material. The microbial communities in the enriched reactors had little resemblance to the original material: e.g. families of the phylum Campylobacterota (*Arcobacteraceae*, *Sulfurimonadaceae*, and *Sulfurospirillaceae*) were abundant in both cysteine (relative abundance between 6% to 62%) and sulfate-enriched reactors (24% to 67%) but were not detected in the original material. In the cysteine-enriched reactors, families *Dethiosulfatibacteraceae*, *Fusobacteriaceae*, and *Vibrionaceae* increased in relative abundance during the experiment, from 31%-55% on day 28 to 67-87% on day 76. In the sulfate-enriched reactors, the relative abundance of families affiliated with phylum Desulfobacterota (*Desulfuromonadaceae*, *Desulfocapsaceae*, *Desulfobacteraceae*, and *Geopsychrobacteraceae*) increased during the experiment, from 7% on day 14 to 27.5% on day 28.

**Figure 2.**
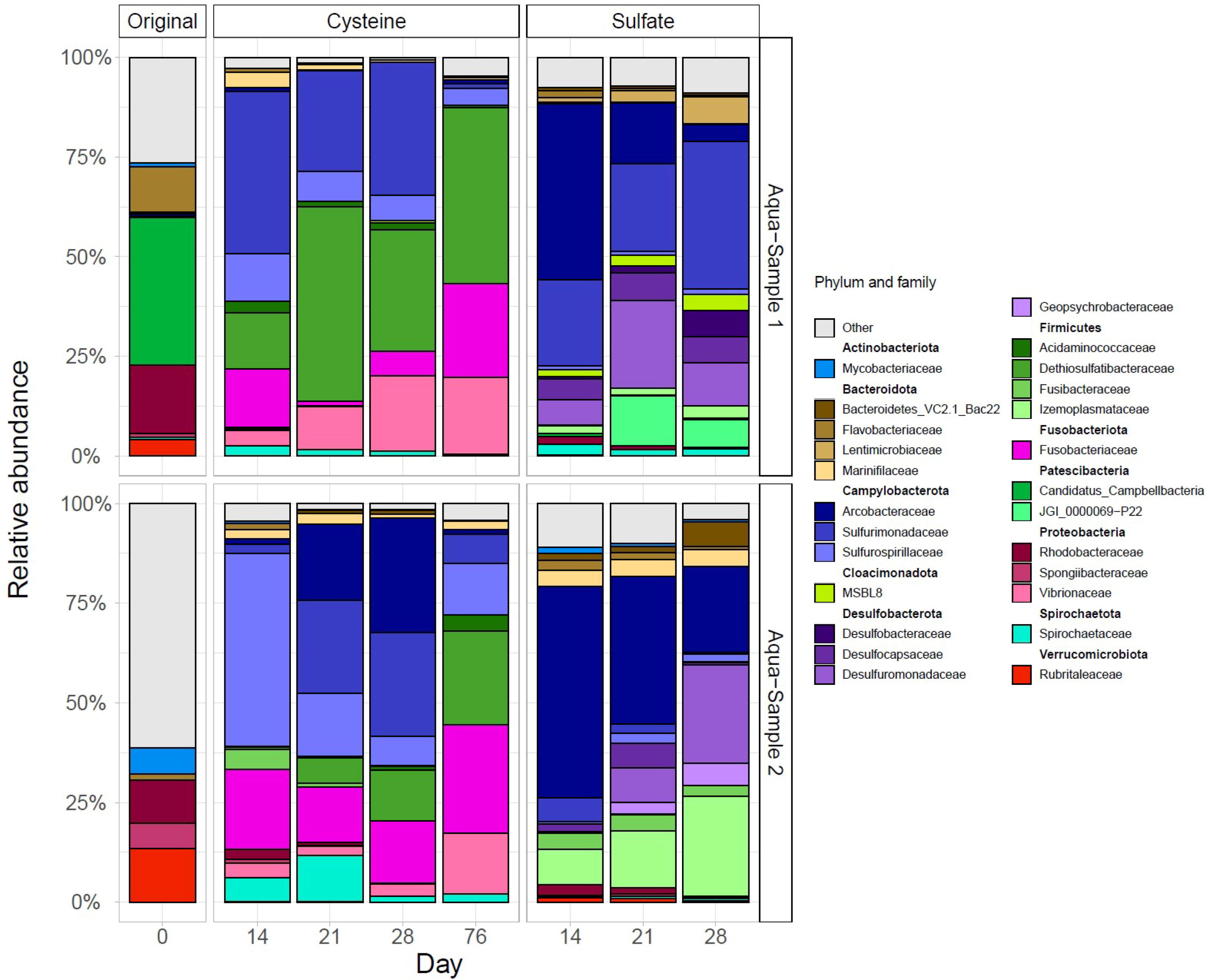
Relative abundances of the top 25 most abundant bacterial families in original material of Aqua-Samples 1 and 2 (raw), at day 14, 21, and 28 under cysteine and sulfate-enriched conditions, and at day 76 under cysteine-enriched condition. “Other” refers to the families outside of the top 25 most abundant ones.

### Metagenomes from enrichment cultures

In order to identify potential bacterial families involved in the production of hydrogen sulfide, we selected a large number of genes that could be involved in the production of hydrogen sulfide via cysteine degradation (*cyuA, mgl, malY, cysK, cysM, sseA, metC, tnaA, dcyD, cbs, cse*), dissimilatory sulfate reduction (*dsrA,B*), and thiosulfate reduction (*phsA/psrA*) (See Supplementary Table S3 for detailed information on selection of these genes). We then examined their abundance in the metagenomes (Fig. 3) and their taxonomic assignment (Fig. 4). We found genes encoding enzymes that may have either cysteine degradation as secondary activity (*mgl, malY, cysK, metC, tnaA, cbs, cse, sseA*) and genes exclusively associated with cysteine degradation (c*yuA, dcyD*) to be present under both sulfate-enriched and cysteine-enriched conditions (Fig. 3). Some of the cysteine-degrading genes (c*ysK, sseA, tnaA, metC*) were found to be present in equal abundances under both conditions, while the abundances of *cbs* (0.03 to 2.14 cpm) and *cse* (0.04 to 2.48 cpm) were very low in all the reactors. Genes *cysM* and *dcyD*, as well as the genes associated with dissimilatory sulfate reduction (*dsrA, B),* were more abundant in the sulfate-enriched than in cysteine-enriched reactors. Three genes associated with cysteine degradation were more abundant in cysteine-enriched reactors: *mgl* (sulfate day 28: 11.15 ± 0.18 cpm, cysteine day 28: 22.45 ± 11.65 cpm, cysteine day 76: 49.59 ± 14.38 cpm), c*yuA* (11.84 ± 2.35 cpm, 37.23 ± 9.14 cpm, 76: 57.61 ± 3.61 cpm) and *malY* (18.70 ± 4.74, 39.4 ± 0.19, 37.3 ± 1.31 cpm). The genes associated with thiosulfate reduction (*phsA*/*psrA*) and oxidation of hydrogen sulfide (*sqr*) were present in all reactors without any significant difference between the treatments.

**Figure 3.**
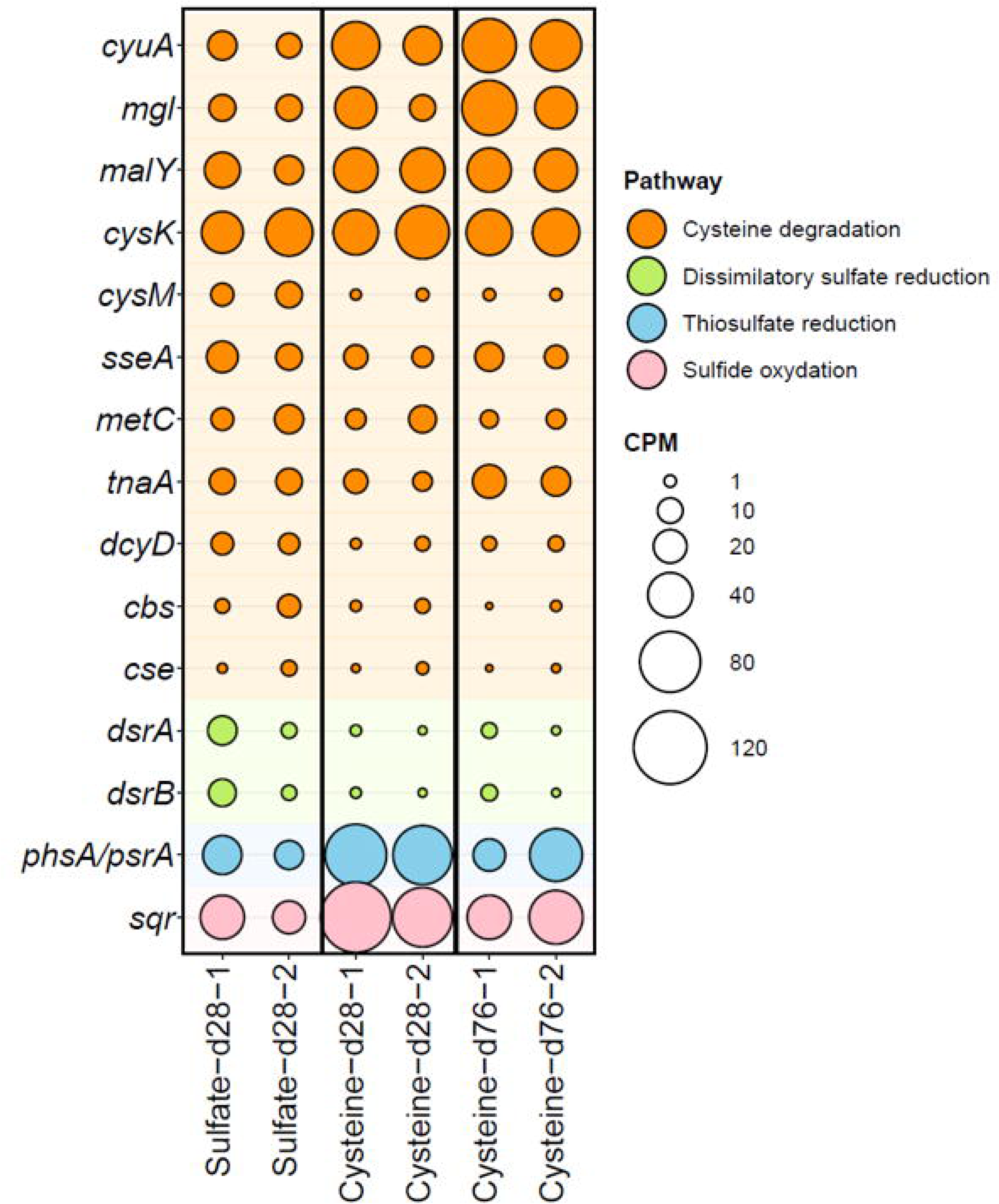
Abundance of genes in the metagenomes reported as coverage per million reads (cpm) with focus of those known to be associated with cysteine degradation in orange, dissimilatory sulfate reduction in green, thiosulfate reduction in blue, and sulfide oxidation gene in pink. Data are for sulfate-enriched reactors at day 28 (Sulfate-d28-1/2) and cysteine-enriched reactors at days 28 and 76 (Cysteine-d28-1/2, Cysteine-d76-1/2).

**Figure 4.**
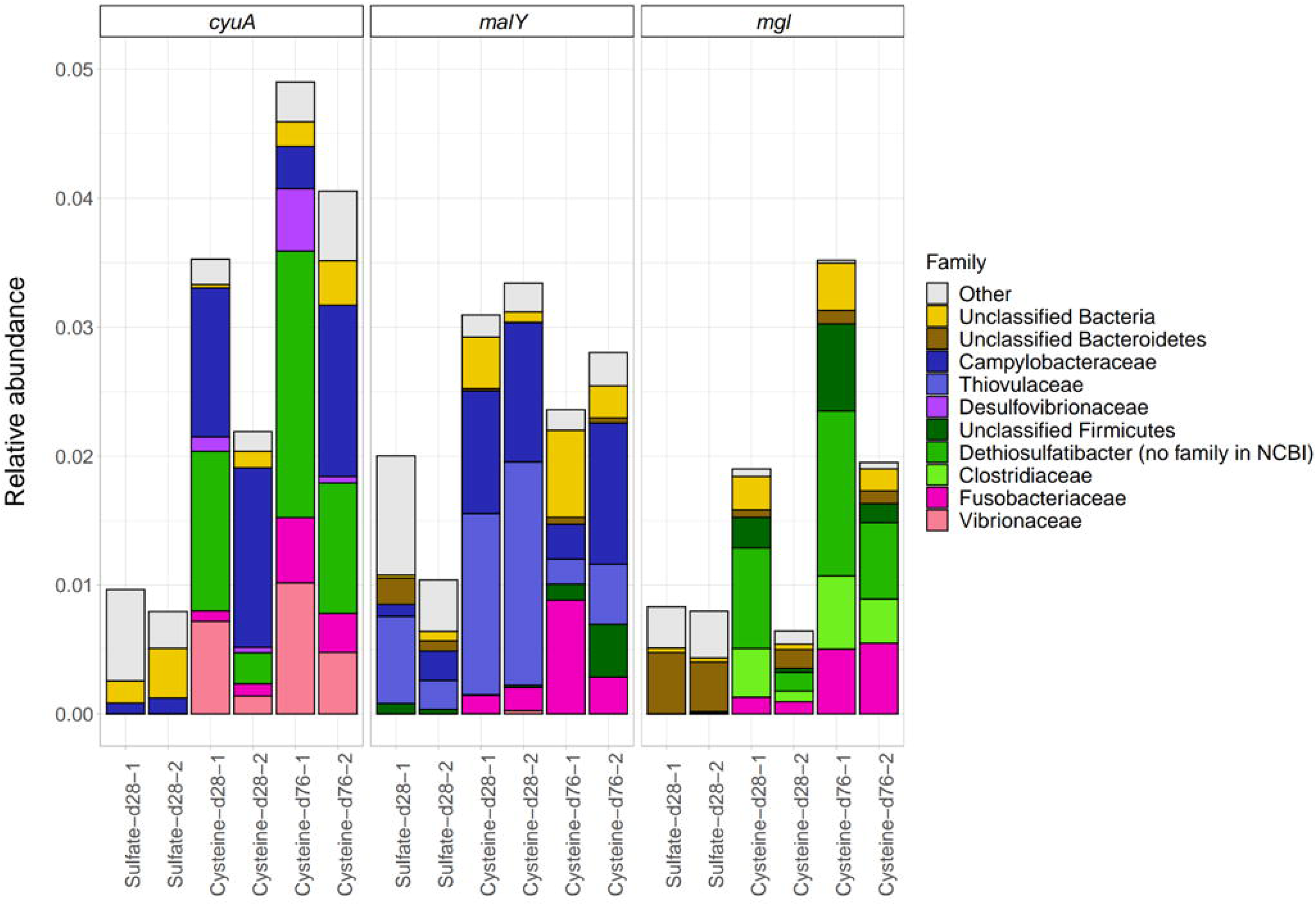
The weight (in percentage of total reads) and taxonomic distribution of *cyuA* (cysteine desulfidase), *malY* (cystathionase), and *mgl* (methionine gamma-lyase) in metagenomes under sulfate-enrichment after 28 days (Sulfate-d28-1/2) and under cysteine-enrichment after 28 and 76 days (Cysteine-d28-1/2, Cysteine-d76-1/2). “Other” refers to the families outside of the 10 most abundant ones.

Further examining the 8 most abundant microbial families carrying the three most abundant H_2_S-producing genes in cysteine-enriched reactors (*cyuA, mgl,* and *malY)* (Fig. 4)*, Fusobacteriaceae* was found to be the only one associated with all three genes. Bacterial families specifically associated with *cyuA* varied according to the enrichment condition. In the sulfate-enriched reactors, the largest share of sequences for c*yuA* was assigned to the families *Campylobacteraceae* and unclassified Bacteria while the rest of the sequences were assigned to families with low relative abundance. In the cysteine-enriched reactors, the largest share of *cyuA* sequences was assigned to the families *Campylobacteraeae*, *Dethiosulfatibacteraceae*, *Vibrionaceae*, *Fusobacteriaceae*, and *Desulfovibrionaceae*. The total abundance of families associated with *malY* was generally higher in cysteine-enriched reactors as compared with sulfate-enriched incubations. In both enrichment conditions, a large portion of the sequences could be assigned to families *Thiovulaceae* and *Campylobacteraceae*. In the cysteine-enriched reactors, gene sequences associated to *Fusobacteriaceae* emerged and alleles associated with families outside the 10 top ones were less abundant than in the sulfate enriched reactors. The total abundance of families associated with *mgl* was generally also higher in cysteine-enriched reactors as compared to sulfate-enriched ones, except for the second cysteine-enriched replicate reactor on day 28. In the sulfate-enriched reactors, most of the gene sequences associated with *mgl* were assigned to unclassified Bacteroidetes and families outside the 10 most abundant ones, while in the cysteine-enriched reactors, the largest share of gene sequences was associated with *Dethiosulfatibacteraceae*, *Clostridiaceae*, *Fusobacteriaceae*, and unclassified Firmicutes.

### Genomes encoding H_2_S-producing pathways

Overall, 71 good quality MAGs (>70% completeness and <10% contamination) and 35 high quality MAGs (>90% completeness and <5% contamination) could be reconstructed from the metagenomic data (Supplementary Fig. S1). MAGs affiliated with *Dethiosulfatibacteraceae*, *Fusobacteriaceae*, and *Vibrionaceae* were selected for in-depth analysis, because of their quantitative increase in cysteine-enriched condition (Fig. 2), but also because of the prevalence of cysteine degrading genes in their genomes (Fig. 4) and the broad representation of these MAGs across the different cysteine reactors (Supplementary Table S4). A MAG associated with *Desulfovibrionaceae* was also included in this pathway mapping, as it was found to be present in cysteine-enriched conditions (Supplementary Fig. S2) and carried the *cyuA* gene (Fig. 4). All four MAGs encode the capacity for at least one cysteine degrading gene (*cyuA, mgl, malY, cysK, sseA, metC, tnaA,* or *dcyD*).

*Dethiosulfatibacteraceae* MAG65b carried five different genes associated with potential cysteine degradation activity (*cyuA, mgl, cysK, sseA,* and *tnaA*). *Psychrilyobacter* MAG139 also carried five different genes with cysteine degradation potential (*cyuA, mgl, malY, cysK,* and *sseA)*, while Vibrio MAG65 carried four genes with cysteine degradation potential (*cyuA, mgl, malY,* and *cysK*). *Desulfovibrionaceae* MAG259 carried all the genes for dissimilatory sulfate reduction (*sat, aprAB,* and *dsrAB*) and four different genes associated with cysteine degradation potential (*cyuA, cysK, metC,* and *dcyD)* (Fig. 5).

**Figure 5.**
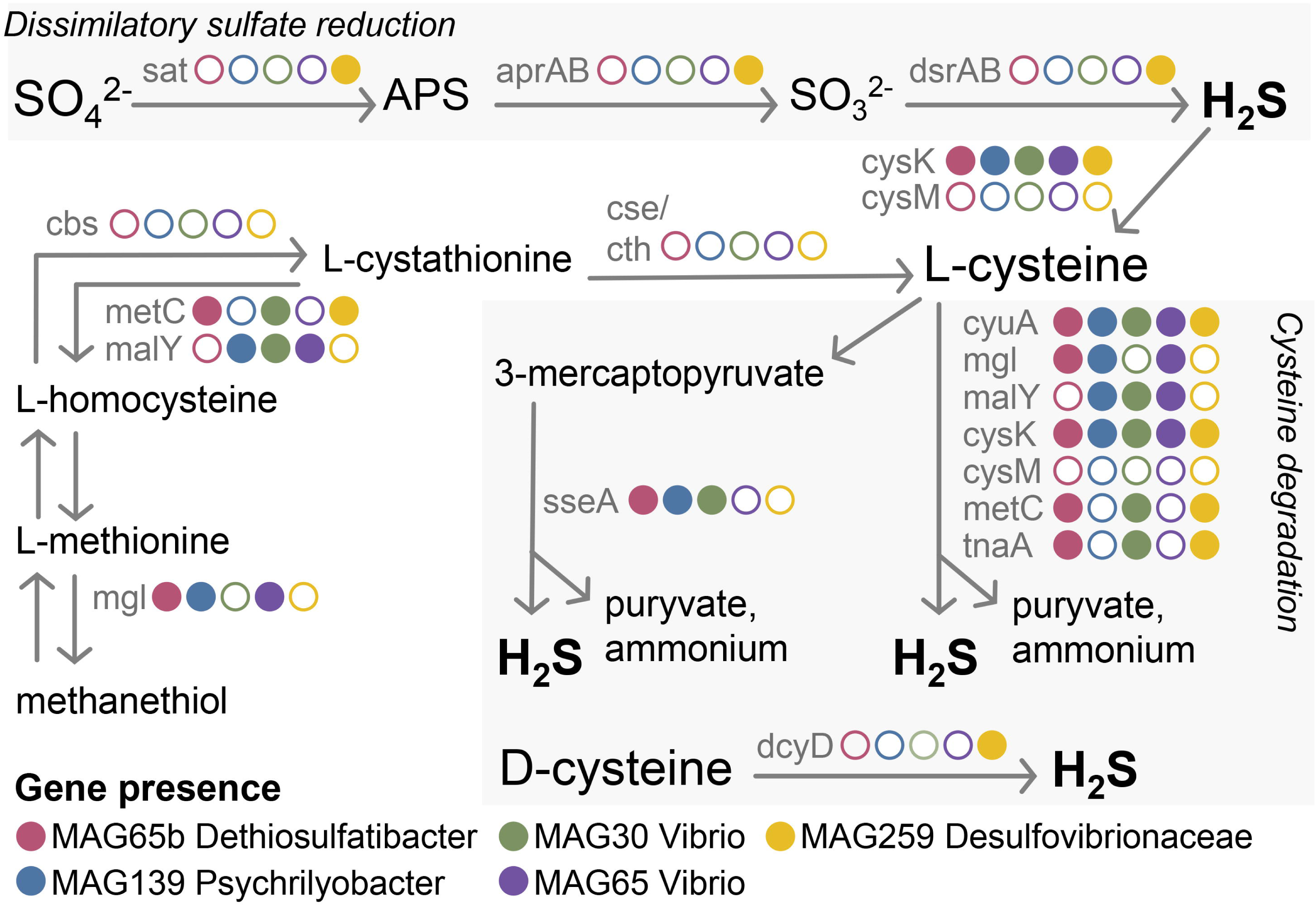
Diagram showing the inferred metabolic pathways responsible for H_2_S production from sulfur and organosulfur compound for the most abundant MAG of the predetermined target families. The presence or absence of the key genes related to the different metabolic features are shown by solid or open circles, respectively, and with different colors for each of the four MAGs. Dissimilatory sulfate reduction pathway is associated with the presence of *sat, aprAB,* and *dsrAB*; cysteine degradation activity is associated with *cyuA, mgl, malY, cysK, cysM, sseA, metC, tnaA, dcyD, cbs,* and *cse*.

### Metagenome assembled genomes

*Dethiosulfatibacteraceae* MAG65b encoded a few amino acid or peptide transporters (Fig. 6A). We detected branched-chain amino acid transporters (*LivKMGF*), the specific cysteine transporter (*cyuP,* also called *yhaO*), and an oligopeptide transporter (*OppABCDF*). In addition to genes involved in cysteine degradation (Fig. 5), MAG65b carried other amino acids degrading enzymes (threonine dehydratase [EC:4.3.1.19], L and D-serine dehydratase [EC:4.3.1.17 4.3.1.18], and alanine dehydrogenase [EC:1.4.1.1]) producing pyruvate and oxobutanoate, which could be fermented to acetate (phosphate acetyltransferase [EC:2.3.1.8], acetate kinase [EC:2.7.2.1], and propionate CoA-transferase [EC:2.8.3.1]), ethanol (alcohol dehydrogenase [EC:1.1.1.1]), or lactate (acryloyl-CoA pathway). The genome also encoded the complete pathway for glycolysis and gluconeogenesis (glucose formation from pyruvate). Overall, we detected 23 peptidase families that could be involved in catabolism of peptides and amino acids.

**Figure 6.**
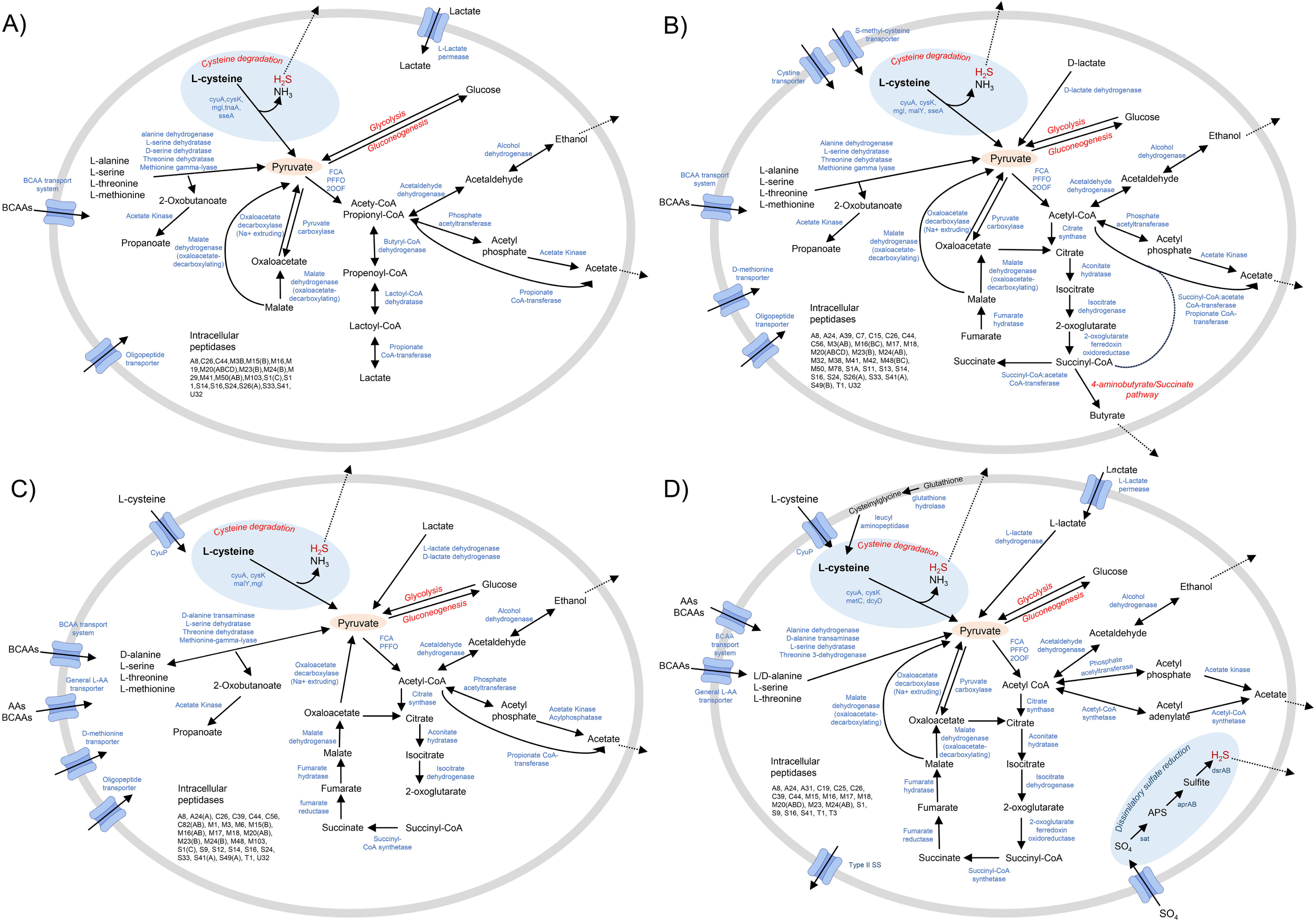
Central metabolism, pyruvate metabolism, and amino acids degradation pathway reconstructed for the four most abundant MAGs associated with the target families and qualified as at least good quality MAGs: A) *Dethiosulfatibacteraceae* MAG65b, (72.4% complete, 3.5% contamination) with no species related from GTDB database, B) *Fusobacteriaceae* MAG139 (99.4% completion, 1.9% contamination) classified as the same species of *Psychrilyobacter* sp002342785 from GTDB database, C) *Vibrionaceae* MAG65 (77.3% complete and 0% contamination) with no species related from GTDB database, and D) *Desulfovibrionaceae* MAG259 (99.4% complete and 9.5% contamination) with no species related from GTDB database. FCA: formate C-acetyltransferase [EC:2.3.1.54]; PFFO: pyruvate-ferredoxin/flavodoxin oxidoreductase [EC:1.2.7.1 1.2.7.-]; 2OOF: 2-oxoglutarate/2-oxoacid ferredoxin oxidoreductase [EC:1.2.7.3 1.2.7.11].

*Fusobacteriaceae* MAG139 encoded several amino acid and peptide transporters (Fig. 6B). We detected an S-methylcysteine transporter (*YxeMNO*), a cystine transporter (*TcyC*), a branched-chain amino acid transporter (*LivK*), a D-methionine transporter (*MetQIN*), and an oligopeptide transporter (*OppABCDF*). In addition to genes involved in cysteine degradation (Fig. 5), MAG139 encoded other amino acid-degrading enzymes (threonine dehydratase [EC:4.3.1.19], L-serine dehydratase [EC:4.3.1.17], and L-alanine dehydrogenase [EC:1.4.1.1]) producing pyruvate and oxobutanoate, which can then be fermented to acetate (phosphate acetyltransferase [EC:2.3.1.8], acetate kinase [EC:2.7.2.1], propionate CoA-transferase [EC:2.8.3.1], and succinyl-CoA:acetate CoA-transferase [EC:2.8.3.18]), ethanol (alcohol dehydrogenase [EC:1.1.1.1]), or lactate (D-lactate dehydrogenase [EC:4.3.1.18]). Produced acetyl-CoA could also feed the TCA cycle (tricarboxylic acid cycle), even if the key enzyme succinate dehydrogenase was missing from the MAG. It could be that the gene was not recruited during the creation of MAG139, although several copies of succinate dehydrogenase associated to *Fusobacteriaceae* family and even an unclassified *Psychrilyobacter* were found in the metagenome samples (Supplementary Data). The genome also encoded the complete glycolysis and gluconeogenesis pathways. Overall, we detected 34 peptidase families that could be involved for catabolism of peptides and amino acids.

*Vibrionaceae* MAG65 encoded several amino acid and peptide transporters (Fig. 6C). We detected the general L-amino acid transporters (*AapJQMP*), ABC-type polar-amino-acid transporter (*ArtIMQP*), D-methionine transporter (*MetQIN*), oligopeptide transporter (*OppABCDF*), and the specific cysteine transporter (*cyuP*). In addition to genes involved in cysteine degradation (Fig. 5), MAG65 encoded other amino acid degrading enzymes (threonine dehydratase [EC:4.3.1.19], L-serine dehydratase [EC:4.3.1.17], and D-alanine transaminase [EC:2.6.1.21]) to produce pyruvate and oxobutanoate which could be further fermented to acetate (phosphate acetyltransferase [EC:2.3.1.8], acetate kinase [EC:2.7.2.1], acylphosphatase [EC:3.6.1.7], and propionate CoA-transferase [EC:2.8.3.1]), ethanol (alcohol dehydrogenase [EC:1.1.1.1]), or lactate (L and D-lactate dehydrogenase [EC:4.3.1.17 4.3.1.18]). Acetyl-CoA produced could also be fed into the TCA cycle, even if 2-oxoglutarate ferredoxin oxidoreductase was missing, probably due to it not being recruited during the creation of the MAG. The genome was also found to encode the complete pathway for glycolysis and gluconeogenesis. Overall, we detected 31 peptidase families that could be involved in catabolism of peptides and amino acids.

*Desulfovibrionaceae* MAG259 encoded different peptide transporters (Fig. 6D). We detected the general L-amino acid transporter (*AapJQMP*), branched-chain amino acid transporter (*LivKHMGF*), L-glutamine transporter (*GlnHPQ*), and the specific cysteine transporter (*cyuP*). In addition to genes involved in cysteine degradation (Fig. 5), MAG259 encoded other amino acid degrading enzymes (threonine 3-dehydrogenase [EC:1.1.1.103], L-serine dehydratase [EC:4.3.1.17], alanine dehydrogenase [EC:1.4.1.1], and D-alanine transaminase [EC:2.6.1.21]) producing pyruvate and oxobutanoate, which could be fermented to acetate (phosphate acetyltransferase [EC:2.3.1.8], acetate kinase [EC:2.7.2.1], and acetyl-CoA synthetase [EC:6.2.1.1]), ethanol (alcohol dehydrogenase [EC:1.1.1.1]) or lactate (L-Lactate dehydrogenase [EC:1.1.1.27]). Acetyl-CoA produced could also feed into the TCA cycle. The genome also encoded the complete pathway for glycolysis, gluconeogenesis, and dissimilatory sulfate reduction. Overall, we detected 21 peptidase families that could be involved for catabolism of peptides and amino acids.

### Gene expression

Of the four selected MAGs, only *Fusobacteriaceae* MAG139 was represented by too few transcripts to enable meaningful expression analyses (Supplementary Table S6). The metatranscriptomes show that in all MAGs*, cyuA* is the only gene expressed for cysteine degradation, and this gene was upregulated significantly and transcribed in all cysteine-fed incubations as compared to sulfate-fed incubations (Fig. 7). In *Dethiosulfatibacteraceae* MAG65b, several genes related to cysteine transport and degradation, pyruvate metabolism, and gluconeogenesis were upregulated when cysteine was available as sulfur source as compared to when sulfate was used as sulfur source (Fig. 7A). Thiosulfate/3-mercaptopyruvate sulfurtransferase ([EC:2.8.1.1 2.8.1.2]), *sseA,* exhibited a substantial TPM increase in response to cysteine, but the expression was not significantly different between sulfate or cysteine-fed incubations. The gene expression for acetate kinase [EC 2.7.2.1] and succinate/propionate CoA-transferase [EC:2.8.3.1], both related to pyruvate metabolism and transformation of acetyl-coA to acetate, also increased significantly in response to cysteine. In contrast, tryptophanase [EC:4.1.99.1], formate C-acetyltransferase [EC:2.3.1.54], pyruvate-ferredoxin/flavodoxin oxidoreductase [EC:1.2.7.1 1.2.7.-], 2-oxoglutarate/2-oxoacid ferredoxin oxidoreductase subunit alpha/beta [EC:1.2.7.3 1.2.7.11], 2,3-bisphosphoglycerate-independent phosphoglycerate mutase [EC:5.4.2.12], phosphoglycerate kinase [EC:2.7.2.3], glyceraldehyde 3-phosphate dehydrogenase [EC:1.2.1.12], and triosephosphate isomerase (TIM) [EC:5.3.1.1] were all significantly upregulated when sulfate was available (Fig. 7A).

**Figure 7.**
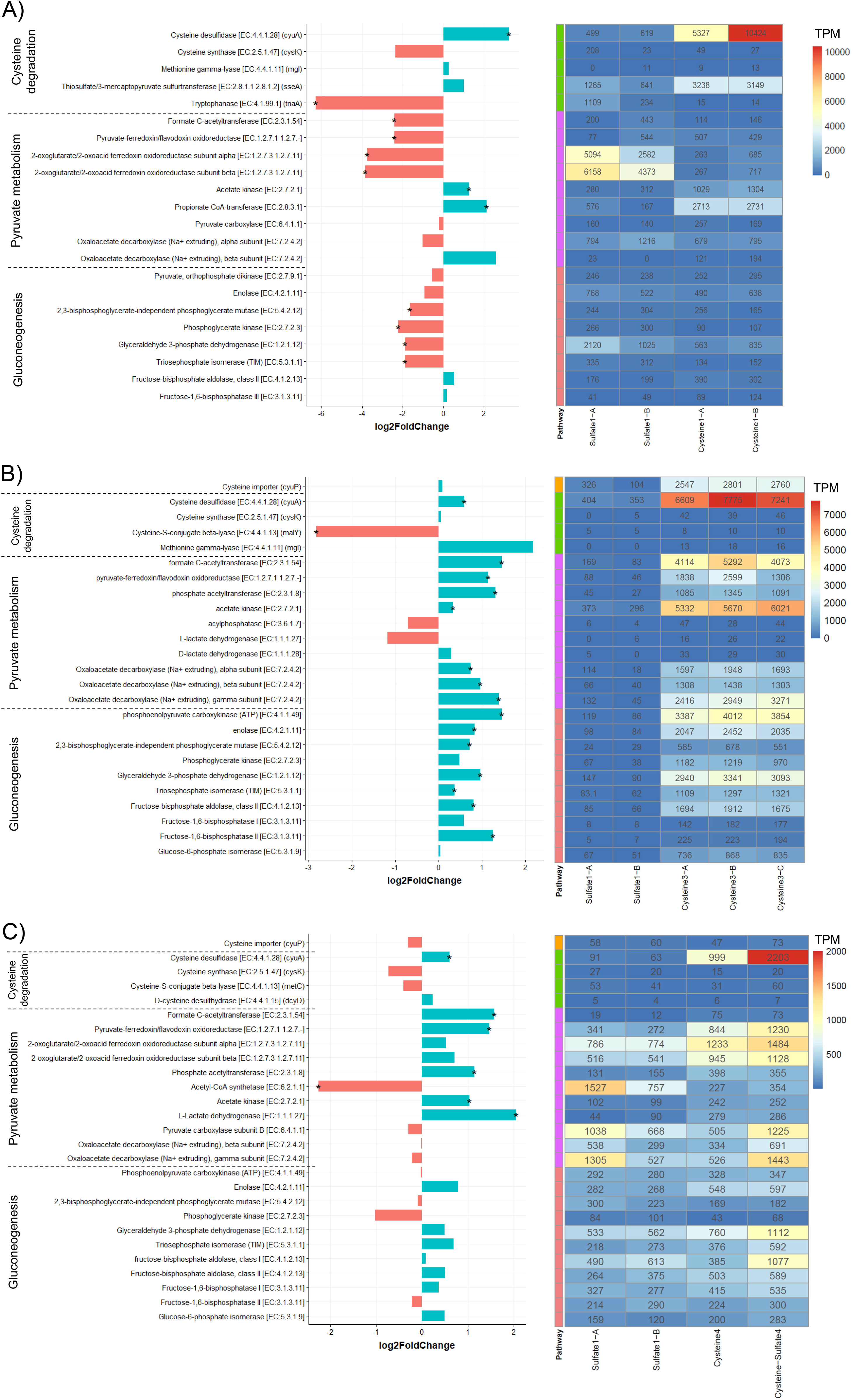
Differences in the gene expression analyzed with DESeq2 and heatmap showing TPM of the genes of interest annotated and grouped according to pathway as cysteine transporters (in yellow), cysteine degradation (in green), pyruvate metabolism in (purple) and gluconeogenesis (in orange) for A) *Dethiosulfatibacteraceae* MAG65b between sulfate or cysteine-fed reactors B) *Vibrionaceae* MAG65 between sulfate or cysteine-fed reactors, and C) *Desulfovibrionaceae* MAG259 between sulfate-fed, cysteine-fed and sulfate+cysteine-fed reactors in experiments 1, 3, and 4. Statistically significant differences (adjusted p < 0.05) in genes expression are marked with an asterisk (*).

In *Vibrio* MAG65, most genes related to cysteine transport and degradation, pyruvate metabolism, and gluconeogenesis were upregulated in response to cysteine as compared to sulfate-fed incubations (Fig. 7B).

However, some upregulated genes exhibited only moderate expression with no significant difference between cysteine and sulfate-fed incubations. For example, *cysK* and *mgl* associated to cysteine degradation and D-lactate dehydrogenase exhibited a low expression in both sulfate and cysteine-fed incubations. The expression of cysteine importer *cyuP*, phosphoglycerate kinase [EC:2.7.2.3], fructose-1,6-bisphosphatase I [EC:3.1.3.11], and glucose-6-phosphate isomerase [EC:5.3.1.9] was higher but not significantly different between cysteine and sulfate-fed incubations. The rest of the genes selected exhibited a significantly higher expression in cysteine-fed incubations as compared to those that were sulfate-fed. Only *malY* (cysteine-S-conjugate beta-lyase [EC:4.4.1.13]) was significantly downregulated in response to cysteine.

In *Desulfovibrionaceae* MAG259, several genes were upregulated when fed cysteine (cysteine only and cysteine and sulfate) as compared to sulfate. Formate C-acetyltransferase [EC:2.3.1.54], pyruvate-ferredoxin/flavodoxin oxidoreductase [EC:1.2.7.1 1.2.7.-], phosphate acetyltransferase [EC:2.3.1.8], acetate kinase [EC:2.7.2.1], and L-lactate dehydrogenase [EC:1.1.1.27] associated with pyruvate metabolism, were significantly upregulated. Acetyl-CoA synthetase [EC:6.2.1.1] was the only gene that was upregulated, with also a clear increase of expression in sulfate-fed incubations. as compared to cysteine-fed incubations. Other genes did not show any clear expression pattern.

## Discussion

In engineered systems, H_2_S often accumulates due to an imbalance between H_2_S production and consumption, causing e.g., sewer corrosion, health hazards, and aquaculture production losses (54,55). So far in RAS, H_2_S has been considered to originate from sulfate reduction, while evidence from natural and engineered ecosystems suggests that microbial degradation of S-rich amino acids can also be an important H_2_S source. In this study, we examined the microbial production of H_2_S in a sulfate-free medium, based exclusively on the S-rich amino acid cysteine and compared the production and microbiology to a “traditional H_2_S source” via the i.e., dissimilatory sulfate reduction pathway.

First and foremost, we could demonstrate microbial H_2_S production from both cysteine and sulfate. The role of cysteine as a significant source of H_2_S was further supported by an increase of acetate and ammonium concentrations in incubation experiments with cysteine. Ammonium is a direct product of the cysteine degradation reaction, originating from the amino group of cysteine released during degradation (56,57). Acetate is not produced directly from cysteine degradation but can be derived from pyruvate produced by cysteine degradation if the bacteria carry the acetate synthesis pathway (58). Conversely, acetate concentration decreased in the incubations with sulfate, attributed to the consumption of acetate by SRM in dissimilatory sulfate reduction (59). We also observed differences in H_2_S production profiles between the two enrichment conditions, as cysteine led to almost immediate H_2_S production, while the process started slower in sulfate-enriched reactors. One putative explanation for this lag could be that the preferred electron acceptors i.e. oxygen, nitrate, and fumarate (60) needed to be fully depleted to allow sulfate reduction. This could also explain why the relative abundance of known SRM (*Desulfuromonadaceae*, *Desulfocapsaceae*, *Desulfobacteraceae*, and *Geopsychrobacteraceae*) was still rather low on day 14 in sulfate-enriched reactors (Fig. 2).

We identified three bacterial families significantly increasing their abundance under cysteine-enriched conditions: *Dethiosulfatibacteraceae*, *Fusobacteriaceae*, and *Vibrionaceae*. Metagenomic analysis showed that members of each of them carried at least one of the three main genes encoding for cysteine degrading enzymes (*cyuA, mgl*, and *malY)* and that the relative abundance of these genes increased in response to cysteine-enrichment. In human gut and oral microbiomes, these same three genes have been identified as the key enzymes/genes for cysteine degradation in diverse bacterial taxa (16,61–64). Our results suggest that these genes and families are the principal ones involved in cysteine based H_2_S production in engineered environments, but potentially also in natural environment, as member of these families have found in lake water column and sediment (65,66), where cysteine degradation activity has also been found (4,67). The rest of our analysis and discussion will be focused on the aforementioned families. We also found a significant share of sequences encoding for one the genes (*cyuA*) to be assigned to family *Desulfovibrionaceae*, and included it in the analysis, even if it was not abundant in the overall microbial community (Fig. 2).

We could successfully reconstruct MAGs for each of the four families: *Dethiosulfatibacteraceae*, *Fusobacteriaceae*, *Vibrionaceae*, and *Desulfovibrionaceae*. *Dethiosulfatibacteraceae* is a bacterial family that is commonly found in marine sediments but has also been observed in engineered environments with potential for H_2_S production (e.g., wastewater treatment or offshore subsea oil storage) (68,69), and in this study, in a recirculating aquaculture system. The members of this family are known to carry the capacity for thiosulfate reduction and could also use a large range of other compounds such as lactate, pyruvate, glutamate, peptone and amino acids like serine, threonine, histidine, lysine, or cysteine for fermentation (70). We could reconstruct two *Dethiosulfatibacteraceae* MAGs (MAG21 and 65b) (Supplementary Fig. S1), but as MAG21 was not abundant nor active, it was not analyzed further (Supplementary Tables S5 and S6). In MAG65b, *cyuA* was the only cysteine degrading gene highly transcribed and upregulated in cysteine-fed reactors in experiments carried out with the cysteine-enriched biomass. As *cyuA* is a cysteine desulfidase, with primary function to degrade cysteine, we suggest this enzyme to be responsible for H_2_S production from cysteine in MAG65b. We also found MAG65b to carry *sseA*, a gene that has two functions: thiosulfate reduction and cysteine degradation. As *Dethiosulfatibacteraceae* is known for capacity to perform thiosulfate-reduction, we expected to find thiosulfate reductase genes (e.g., thiosulfate reductase (quinone) genes (phsABC)) but did not detect any such gene other than sseA. In our experiments, we detected an increase of sseA transcription in cysteine-fed reactors, but this increase was not significant when compared to sulfate-fed reactors. It is still possible that thiosulfate was produced and then reduced to H_2_S by *sseA* in our reactors, which would explain why this gene was transcribed under both cysteine and sulfate amended conditions. More in-depth functional studies will need to be carried out for *Dethiosulfatibacteraceae* in order to understand how *sseA* reacts and how it is involved in cysteine and thiosulfate metabolism. Our metagenome analysis confirmed that our MAG65b *Dethiosulfatibacteraceae* could use cysteine either for gluconeogenesis or fermentation of pyruvate to acetate, as all pathways and key enzymes were present (Fig. 6A). Moreover, metatranscriptome analysis revealed acetate kinase and succinate/propionate CoA-transferase to be upregulated with a relatively high transcription in cysteine-fed compared to sulfate-fed reactors (Fig. 7A), indicating that cysteine has been mostly used for fermentation purposes and ATP production (71,72). This acetate production explains the observed increase of acetate concentration in the 48h-experiment when cysteine was added (Table 1).

Unlike *Dethiosulfatibacteraceae*, members of the family *Fusobacteriaceae* (genus *Fusobacterium*) have been shown to produce H_2_S via the cysteine degradation pathway in human bodies (12,28,29), with methionine gamma-lyase (mgl) as a key enzyme (16,61). The presence of *Fusobacterium* has also been found to coincide with H_2_S production from aquaculture organic waste (26). In our cysteine-enriched samples, the key cysteine degrading genes sequences were all associated with the *Fusobacteriaceae* family (Fig. 4), and most of these gene sequences were annotated to the *Fusobacterium* genus (see Supplementary Data). These findings highlight the importance of this family for non-sulfate based H_2_S production in the presence of S-rich amino acids and absence of oxygen. Unfortunately, we could not reconstruct any good quality MAGs associated with *Fusobacterium*. However, we did reconstruct a *Fusobacteriaceae* MAG139 with 96% identity to *Psychrilyobacter sp02342785* (GTDB). Previously, the *Psychrilyobacter* genus has been found to use a wide range of amino acids and peptides for growth in both marine sediments (73) and abalone gut (74). Indeed, we found this MAG to carry an array of peptidases, different transporters for either oligopeptide or amino acids, and different genes for the fermentation of amino acids. It also carried four genes involved in cysteine degradation. Although we did not obtain enough metatranscriptomic reads from the *Psychrilyobacter* MAG to enable further analysis (Supplementary Tables S5 and S6), the high abundance of *Fusobacteriaceae* in the cysteine-enriched reactors and the fact that they harbor key cysteine degrading genes combined with earlier findings in the medical field, strongly suggests that *Psychrilyobacter* and/or the broader *Fusobacteriaceae* family play an important role in the production of H_2_S from cysteine also in engineered ecosystems.

The family *Vibrionaceae* is mainly known for its many pathogens capable of causing human and animal disease (75,76), but this diverse genus also includes other members native to a wide range of aquatic habitats (e.g., sediment, column water, deep sea, or as symbionts (65,77)). The *Vibrionaceae* family is known to be capable of cysteine degradation (78), and at least some members, such as *Vibrio cholerae,* are using the putative cystathionine B-synthase (*cbs*) for H_2_S-producing cysteine degradation (79). We did not detect any cbs genes in the metagenomes, but found the genes *cyuA, malY, cysK, sseA, metC*, and *tnaA* associated with the *Vibrionaceae* family. We could reconstruct two MAGs (MAG30, 65) associated to the genus *Vibrio*. MAG30 carried all the cysteine degrading genes listed above but was not abundant and showed little presence and expression in the metatranscriptomes, suggesting it not being active (Supplementary Tables S5 and S6). In contrast, the second *Vibrio* MAG (MAG65), containing the genes *cyuA, mgl, malY,* and *cysK,* appeared to be extremely active based on our community-wide expression profiling. The *cyuPA* (also known as *yhaOM*) operon is known to be the major anaerobic cysteine transporter and cysteine desulfidase in *Escherichia coli* and *Salmonella enterica* (63). We found these genes to be highly expressed in our study (Fig. 7B), but only *cyuA* was upregulated significantly in cysteine-fed reactors as compared to sulfate-fed reactors, suggesting it to be the main enzyme responsible for cysteine degradation and H_2_S production in our *Vibrio* MAG65. To the best of our knowledge, this is the first time that *cyuA* has been identified as the key gene for H_2_S production in a *Vibrio sp*. Previously, some members of the *Vibrio* genus have been found to ferment glucose to pyruvate and subsequently transform this metabolite further to acetate (80). We found Vibrio MAG65 to carry such a complete pathway for pyruvate fermentation to acetate and genes related to this fermentation were all upregulated and highly transcribed in cysteine-fed reactors compared to sulfate-fed reactors (Fig. 7B). This inferred acetate production can explain the observed increase of acetate concentration in the 48h-experiment when cysteine was added (Table 1). We also observed the transcription of all genes for gluconeogenesis, a majority being significantly upregulated in cysteine-fed reactors as compared to sulfate-fed treatments, suggesting that pyruvate produced from cysteine was also transformed into glucose as an energy-reserve and that cysteine was not only used for sulfur and ATP but also as a carbon source in our Vibrio MAG65. In addition to the three above-mentioned families, we also found the family *Desulfovibrionaceae* to be of interest. *Desulfovibrionaceae* is a well-known SRM, which is commonly found in marine environments but also in engineered H_2_S-rich environments (e.g., aquaculture, wastewater) (6,81–83). However, we did not detect the *Desulfovibrionaceae* family in the sulfate-enriched reactors but only in cysteine-enriched reactors (Supplementary Fig. S2). This could be explained by the media used in this study. The media contained a combination of formate, acetate, and propionate, as it was formulated following previous knowledge on optimal SRM enrichment media. However, some genera within the *Desulfovibrionaceae* lack the capacity to use acetate or propionate to fuel dissimilatory sulfate reduction (84,85), meaning that they would be restricted to use formate for this purpose. Hence when formate would have been depleted in our reactors, *Desulfovibrionaceae* would not be able to survive in sulfate media, while in cysteine-enriched media, they could potentially have used cysteine as an alternative carbon source for fermentation and growth, as previously suggested for *Desulfovibrionaceae* (86,87). Indeed, our metagenome and metatranscriptome analysis confirm the potential of *Desulfovibrionaceae* MAG259 to function as a cysteine degrader and fermenter in addition to being a sulfate reducing bacteria. Of the four cysteine degrading genes seen in this population, only *cyuA* was highly transcribed and upregulated in cysteine-fed reactors compared to sulfate-fed reactors (Fig. 7C), strongly suggesting that *cyuA* was also the key enzyme for cysteine degradation in our *Desulfovibrionaceae* MAG259. Pyruvate produced by *cyuA* could have been converted into acetyl-coA and then acetate for energy acquisition and ATP production explaining the upregulation of formate C-acetyltransferase, pyruvate-ferredoxin/flavodoxin oxidoreductase, phosphate acetyltransferase, and acetate kinase in cysteine-fed incubations as compared to sulfate-fed ones. Analogous to the previous MAGs, this acetate production can explain observed increases in acetate concentration during the 48h-experiment when cysteine was added (Table 1). Pyruvate could also be used by *Desulfovibrionaceae* MAG259 to produce lactate or formate by either L-lactate dehydrogenase or formate C-acetyltransferase. Both lactate and formate could then be used for dissimilatory sulfate reduction if sulfate is available (84,85). Finally, pyruvate could also be transformed into glucose via gluconeogenesis, as described for the previous MAG. Therefore, in our MAG259, cysteine could have a central place in the metabolism of the cell as either a carbon or energy source or as a source of organic matter to drive dissimilatory sulfate reduction.

By combining production measurements, amplicon sequencing, and metagenomic and metatranscriptomic analysis, we demonstrated that cysteine degradation can be an important H_2_S source with four microbial families centrally involved in this process. *CyuA* seems to be the main gene responsible for cysteine degradation, and while there is previous evidence on the ability of two of these families (*Fusobacteriaceae* and *Vibrionaceae*) to carry out H_2_S production through cysteine degradation, we also identified one new family (*Dethiosulfatibacteraceae*) to be involved in H_2_S production from cysteine and one family (*Desulfovibrionaceae*) to have the potential to engage in either SRM or cysteine degradation. We also showed that cysteine can be used as a carbon and energy source by some bacteria to fuel general cellular anabolic and catabolic needs.

While in some engineered systems, the supply and inputs of cysteine and other S-rich amino acids is limited, environments such as sewage, biogas facilities, and aquaculture systems are by design rich in proteins, enabling and enhancing non-sulfate based H_2_S production. In line with this, a major share of H_2_S has been found to originate from S-rich amino acids in anaerobic digesters and H_S_S production has been found to increase with sludge protein content in aquaculture systems (25,26). Our results corroborate these previous findings but also provide new knowledge on the identity and metabolism of the microbial groups involved in non-sulfate based H_2_S production in protein-rich engineered environments, such as aquaculture systems. Overall, our findings encourage further research on H_2_S and sulfur cycling in aquaculture environments, especially in freshwater aquaculture systems, where sulfate concentrations are low while non-sulfate based H_2_S production can still be fueled by the accumulating organic matter.

## Acknowledgements

We thank Ulla Sproegel, Brian Moller, and Signe Østergaard (DTU AQUA) for their help and expertise in water analysis. We also thank Lauren Davies and Prune Leroy (SLU) for their help in the laboratory for DNA/RNA extraction and sequencing.

## Author contributions

AN and SLA designed the study with input from SB. AN carried out the experimental work and chemical analysis with support from SLA. AN performed amplicon sequencing and prepared samples for metagenomic and metatranscriptomic analysis with support from SB. AN and FPS performed bioinformatics. All authors contributed to data interpretation. AN prepared the first version of the manuscript with major contributions from SLA and additional inputs from FPS and SB in shaping the final manuscript.

## Supplementary material

Supplementary material is available at The ISME Journal online.

## Data availability

Amplicon sequencing data sets, metagenome-assembled genomes, and metagenome and metatranscriptome reads are all available under NCBI-Genbank Bioproject PRJNA1083305.

