## Supplementary data for "Identification of bacteria involved in non-sulfate based hydrogen sulfide production in an aquaculture environment"

**Non-sulfate-based hydrogen sulfide production in engineered systems**

Alexandre Nguyen-tiêt[1], Fernando Puente-Sánchez[2], Stefan Bertilsson[2], Sanni L. Aalto[1]
[1]Technical University of Denmark, DTU Aqua, Section for Aquaculture, Hirtshals, Denmark
[2]Swedish University of Agricultural Sciences, Department of Aquatic Sciences and Assessment, Uppsala, Sweden

### Supplementary Figures

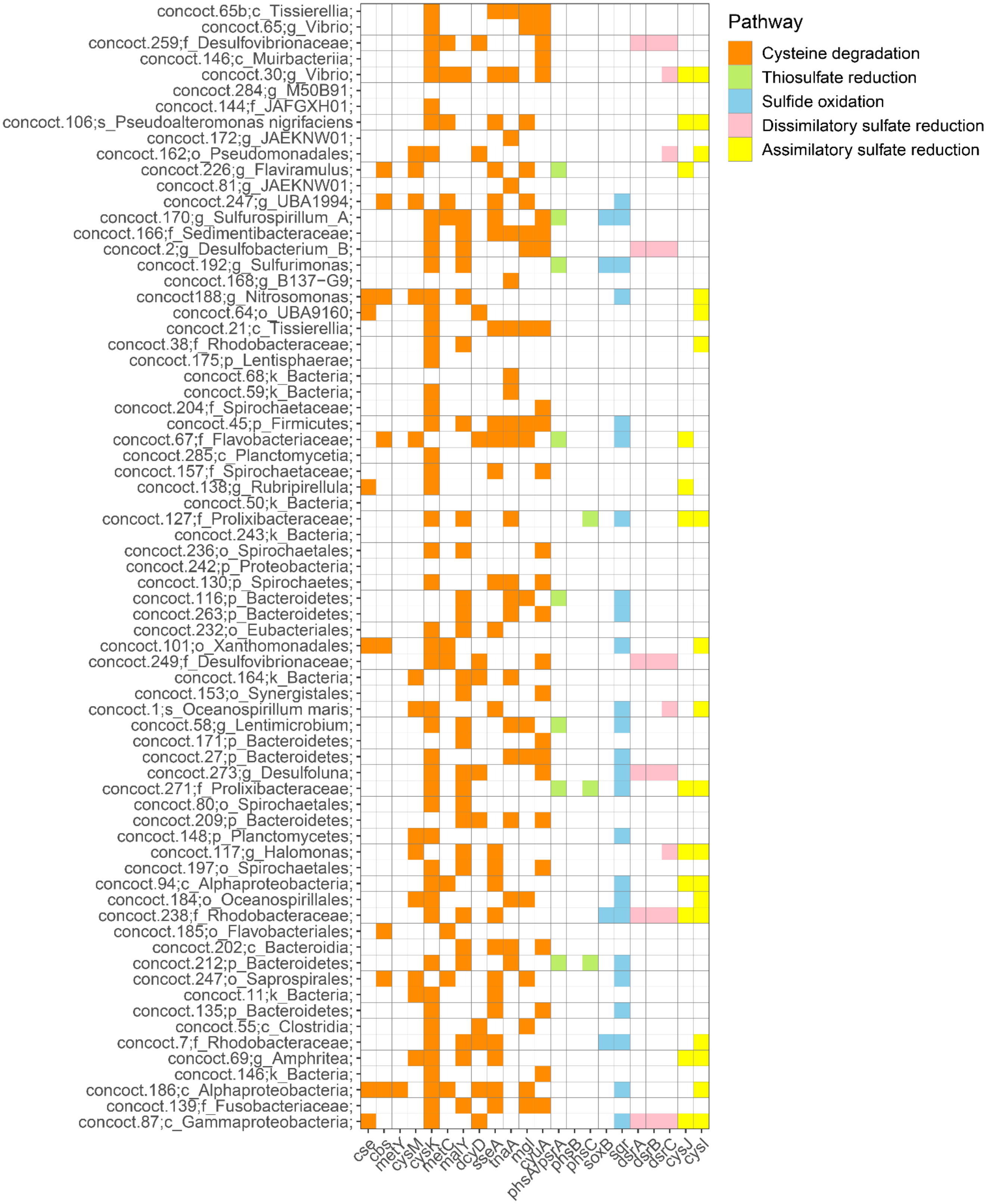


**Figure S1.** Presence and absence of the genes of interest for all MAGs considered as at least “good quality MAGs (>70% completeness and <10% contamination)”.

**Non-sulfate-based hydrogen sulfide production in engineered systems**
Alexandre Nguyen-tiêt[1], Fernando Puente-Sánchez[2], Stefan Bertilsson[2], Sanni L. Aalto[1]
[1]Technical University of Denmark, DTU Aqua, Section for Aquaculture, Hirtshals, Denmark
[2]Swedish University of Agricultural Sciences, Department of Aquatic Sciences and Assessment, Uppsala, Sweden

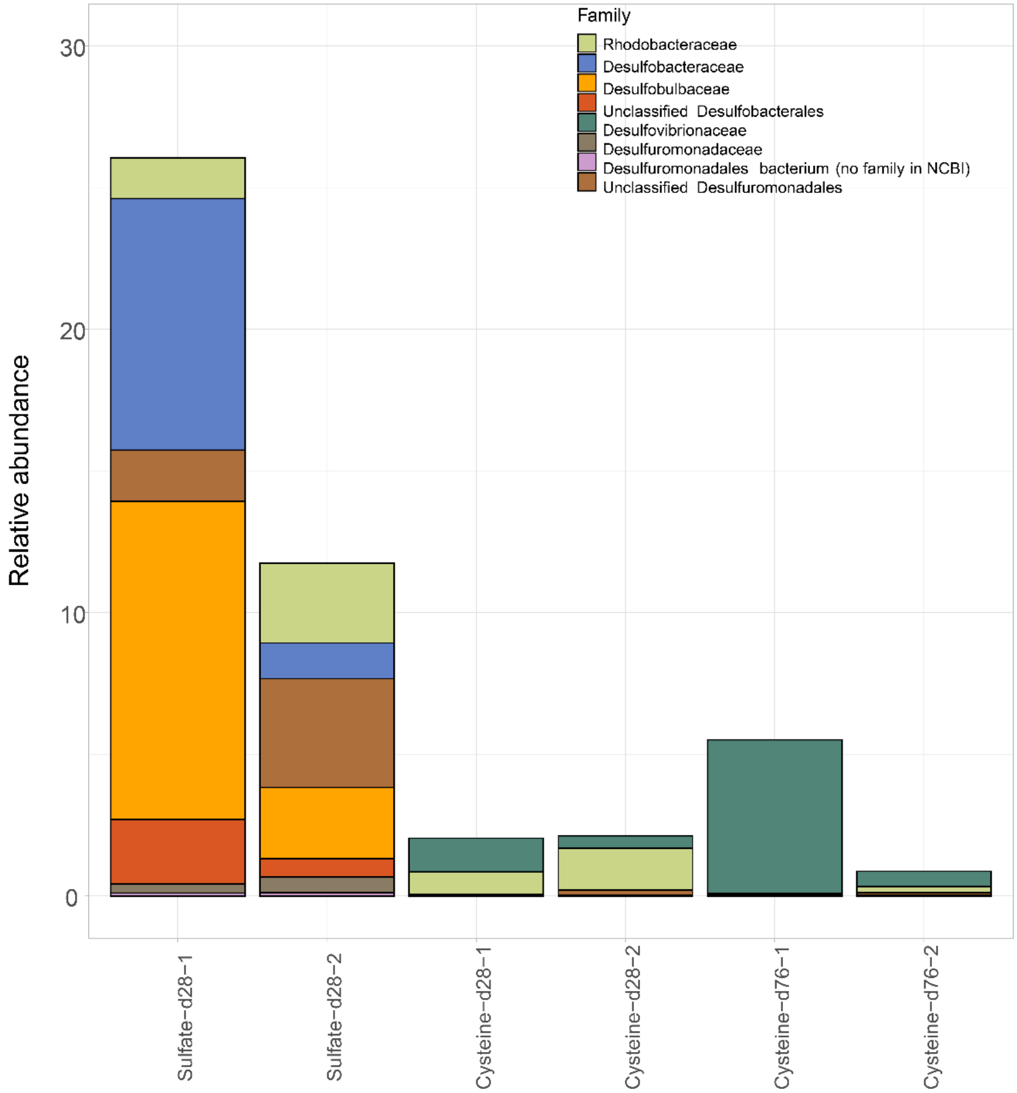


**Figure S2.** Relative abundance of families associated with sulfate reducing bacteria in metagenomes under sulfate-enrichment after 28 days (Sulfate-d28-1/2) and under cysteine-enrichment after 28 and 76 days (Cysteine-d28-1/2, Cysteine-d76-1/2).
