## Supplementary material for "Identification of bacteria involved in non-sulfate based hydrogen sulfide production in an aquaculture environment"

**Non-sulfate-based hydrogen sulfide production in engineered systems**

Alexandre Nguyen-tiêt, Fernando Puente-Sánchez, Stefan Bertilsson, Sanni L. Aalto

**Supplementary Table S1.** Culture media recipe for enrichment reactors and experiments 1-4.

| **Receipt** | **ml** |
|---|---|
| *Solution A* | 890 |
| *Solution B* | 30 |
| *Solution C* | 10 |
| *Solution D* | 50 |
| Autoclave solution A in 1 L bottle. To complete medium, add appropriate amounts of solutions B to D to A in the sequence as indicated. The final pH of the complete medium should be 7.0 - 7.4. | |

| **Solution A** | | |
|---|---|---|
| Chemical | Amount | Unit |
| *$KH_2PO_4$* | *0.2* | *g* |
| *$NH_4Cl$* | *0.3* | *g* |
| *NaCl* | *28* | *g* |
| *$MgCl_2$ x 6 $H_2O$* | *5.5* | *g* |
| *KCl* | *0.7* | *g* |
| *$CaCl_2$ x 2 $H_2O$* | *1* | *g* |
| *Na-resazurin (0.1% w/v)* | *0.5* | *mL* |
| *Selenite-tungstate solution [1]* | *1* | *mL* |
| *Trace element solution SL10 [2]* | *1* | *mL* |
| *Only for sulfate media: $Na_2SO_4$** | *3* | *g* |
| *Only for cysteine media: L-cysteine** | *3* | *g* |
| *Distilled water* | *890* | *mL* |
| **Both sulfur sources added in media used in experiment 4* | | |

| **Solution B** | | |
|---|---|---|
| Chemical | Amount | Unit |
| $Na_2CO_3$ | 1.5 | g |
| *Distilled water* | 30 | ml |

| **Solution C** | | |
|---|---|---|
| Chemical | Amount | Unit |
| *Na-propionate* | 1 | g |
| *Na-acetate* | 1 | g |
| *Na-formate* | 0.5 | g |
| *Distilled water* | 50 | ml |

| **Solution D** | | |
|---|---|---|
| Chemical | Amount | Unit |
| Biotin | 2 | mg |
| Folic acid | 2 | mg |
| Pyridoxine hydrochloride | 10 | mg |
| Thiamine-HCl | 5 | mg |
| Riboflavin | 5 | mg |
| Nicotinic acid | 5 | mg |
| D-Ca-pantothenate | 5 | mg |
| Vitamin B12 | 0.1 | mg |
| p-aminobenzoic acid | 5 | mg |
| Lipoic acid | 5 | mg |
| Distilled water | 1000 | ml |

**Supplementary Table S2.** Concentration and production rates (as mg/l/d) of four VFAs (formate, propionate, butyrate, and valerate) during 48h experiments in reactors with cysteine-enriched biomass fed with either cysteine or sulfate media. Production rate with minus means that the component was consumed, not produced.

| Time (hours) | 0 | 48 | Production rate mg/l/d |
|---|---|---|---|
| **Formate (mg/l)** | | | |
| Aqua-Sample-2 with cysteine (n=3) | 282 ± 1.27 | 4.40 ± 3.63 | -139 ± 1.19 |
| Aqua-Sample-2 with sulfate (n=3) | 274 ± 1.68 | 0 ± 0.00 | -137 ± 0.84 |
| **Propionate (mg/l)** | | | |
| Aqua-Sample-2 with cysteine (n=3) | 742 ± 3.38 | 744 ± 29.2 | 1.00 ± 12.9 |
| Aqua-Sample-2 with sulfate (n=3) | 750 ± 0.73 | 757± 1.32 | 3.68 ± 1.03 |
| **Butyrate (mg/l)** | | | |
| Aqua-Sample-2 with cysteine (n=3) | nd | nd | nd |
| Aqua-Sample-2 with sulfate (n=3) | nd | nd | nd |
| **Valerate (mg/l)** | | | |
| Aqua-Sample-2 with cysteine (n=3) | nd | nd | nd |
| Aqua-Sample-2 with sulfate (n=3) | nd | nd | nd |

**Supplementary Table S3.** The abbreviation, name, code, database used, primary and possible secondary functions, and references of the genes that could be involved in the production of hydrogen sulfide via cysteine degradation, dissimilatory sulfate reduction, and thiosulfate reduction used in this study.

| Abbreviation | Name | Code | Database | Primary function | Secondary function | Reference |
|---|---|---|---|---|---|---|
| cyuA (yhaM) | Putative L-cysteine desulfidase CyuA | COG3681 | EggNOG | L-cysteine desulfidase | None | (3) |
| mgl | Methionine gamma-lyase | K01761 | Kegg | L-methionine methanethiol-lyase (deaminating; 2-oxobutanoate-forming) | Cysteine desulfhydrase | (4) |
| malY | Cysteine-S-conjugate beta-lyase | K14155 | Kegg | L-cysteine-S-conjugate thiol-lyase | Cysteine desulfhydrase | (5) |
| cysK | Cysteine synthase A | K01738 | Kegg | Cysteine synthase | Cysteine desulfhydrase | (5) |
| cysM | S-sulfo-L-cysteine synthase (O-acetyl-L-serine-dependent) | K12339 | Kegg | Cysteine synthase | Cysteine desulfhydrase | (5) |
| sseA (3MST) | Thiosulfate/3-mercaptopyruvate sulfurtransferase | K01011 | Kegg | Sulfurtransferase | | (6) |
| metC | Cysteine-S-conjugate beta-lyase | K01760 | Kegg | L-cysteine-S-conjugate thiol-lyase | Cysteine desulfhydrase | (5) |
| tnaA | Tryptophanase | K01667 | Kegg | Tryptophanase | Cysteine desulfhydrase | (5) |
| dcyD | D-cysteine desulfhydrase | K05396 | Kegg | D-cysteine desulfhydrase | | (7) |
| cbs | Cysteine-S-conjugate beta-lyase | K01697 | Kegg | L-serine hydro-lyase | Cysteine desulfhydrase | (8) |
| cse | Cystathionine-gamma-lyase | K01758 | Kegg | Cystathionase | Cysteine desulfhydrase | (8) |
| dsrA | Dissimilatory sulfite reductase alpha subunit | K11180 | Kegg | Dissimilatory sulfite reductase | | (9) |
| dsrB | Dissimilatory sulfite reductase beta subunit | K11181 | Kegg | Dissimilatory sulfite reductase | | (9) |
| phsA/psrA | Thiosulfate reductase / polysulfide reductase chain A | K08352 | Kegg | Thiosulfate---dithiol sulfurtransferase | | (10) |
| sqr | Sulfide:quinone reductase | K17218 | Kegg | Sulfide:quinone oxidoreductase | | (11) |

**Supplementary Table S4.** The relative abundance of good quality MAGs associated with Tissierellia, Fusobacteriaceae, Vibrionaceae, and Desulfovibrionaceae in metagenomes under sulfate-enrichment after 28 days (Sulfate-d28-1/2) and under cysteine-enrichment after 28 and 76 days (Cysteine-d28-1/2, Cys-d76-1/2).

| MAG | Sulfate-d28-1 | Sulfate-d28-2 | Cysteine-d28-1 | Cysteine-d28-2 | Cysteine-d76-1 | Cysteine-d76-2 |
|---|---|---|---|---|---|---|
| MAG21 (*Tissierellia*) | 0.001% | 0.004% | 0.03% | 0.03% | 9.04% | 0.15% |
| MAG65b (*Dethiosulfatibacter*) | 0.002% | 0.003% | 9.29% | 1.75% | 15.1% | 7.42% |
| MAG139 (*Psychrilyobacter*) | 0.003% | 0.004% | 0.03% | 0.24% | 0.006% | 0.03% |
| MAG30 (*Vibrio*) | 0.001% | 0.002% | 1.35% | 1.52% | 1.83% | 0.85% |
| MAG65 (*Vibrio*) | 0.002% | 0.001% | 14.3% | 2.09% | 19.5% | 9.44% |
| MAG249 (*Desulfovibrionaceae*) | 0.009% | 0.003% | 0.46% | 0.15% | 0.13% | 0.002% |
| MAG259 (*Desulfovibrionaceae)* | 0.002% | <0.001% | 0.78% | 0.22% | 5.91% | 0.56% |

**Supplementary Table S5.** The relative abundance of good quality MAGs associated with Tissierellia, Fusobacteriaceae, Vibrionaceae, and Desulfovibrionaceae in metatranscriptomes collected in experiments 1, 3, and 4.

| MAG | Sulfate1-A | Sulfate1-B | Cysteine1-A | Cysteine 1-B |
|---|---|---|---|---|
| MAG21 (Tissierellia) | 0.006% | 0.01% | 0.11% | 0.05% |
| MAG65b (Dethiosulfatibacter) | 0.36% | 0.97% | 5.20% | 8.33% |
| MAG139 (Psychrilyobacter) | 0.04% | 0.004% | 0.01% | 0.26% |

| MAG | Sulfate1-A | Sulfate1-B | Cysteine3-A | Cysteine3-B | Cysteine3-C |
|---|---|---|---|---|---|
| MAG30 (Vibrio) | 0.07% | 0.03% | 0.43% | 0.65% | 0.63% |
| MAG65 (Vibrio) | 0.36% | 0.97% | 5.2% | 8.33% | 7.44% |

| MAG | Sulfate1-A | Sulfate1-B | Cysteine4 | Cysteine +Sulfate4 |
|---|---|---|---|---|
| MAG249 (*Desulfovibrionaceae*) | 13.6% | 0.48% | 30.7% | 45.2% |
| MAG259 (*Desulfovibrionaceae)* | 0.92% | 0.24% | 0.46% | 0.48% |

**Supplementary Table S6**. Abundance of genes associated with cysteine degradation in MAGs classified either at Tissierellia, Fusobacteriaceae, Vibrionaceae, and Desulfovibrionaceae in metatranscriptomes collected in experiments 1, 3, and 4.

| MAG | Gene | Sulfate1-A | Sulfate1-B | Cysteine1-A | Cysteine1-B |
|---|---|---|---|---|---|
| MAG21 (*Tissierellia*) | *cyuA* | 4 | 2 | 118 | 2 |
| MAG21 (*Tissierellia*) | *mgl* | 0 | 0 | 42 | 1 |
| MAG21 (*Tissierellia*) | *cysK* | 0 | 0 | 0 | 0 |
| MAG21 (*Tissierellia*) | *sseA* | 0 | 0 | 92 | 115 |
| MAG21 (*Tissierellia*) | *tnaA* | 0 | 0 | 0 | 0 |
| MAG65b (*Dethiosulfatibacter*) | *cyuA* | 119 | 118 | 22588 | 34980 |
| MAG65b (*Dethiosulfatibacter*) | *mgl* | 0 | 0 | 33 | 40 |
| MAG65b (*Dethiosulfatibacter*) | *cysK* | 34 | 3 | 147 | 63 |
| MAG65b (*Dethiosulfatibacter*) | *sseA* | 214 | 86 | 9972 | 7682 |
| MAG65b (*Dethiosulfatibacter*) | *tnaA* | 277 | 47 | 68 | 52 |
| MAG139 (*Psychrilyobacter*) | *cyuA* | 6 | 0 | 6 | 350 |
| MAG139 (*Psychrilyobacter*) | *mgl* | 51 | 0 | 8 | 18 |
| MAG139 (*Psychrilyobacter*) | *malY* | 18 | 0 | 0 | 459 |
| MAG139 (*Psychrilyobacter*) | *cysK* | 0 | 0 | 0 | 61 |
| MAG139 (*Psychrilyobacter*) | *sseA* | 0 | 0 | 0 | 9 |

| MAG | Gene | Sulfate1-A | Sulfate1-B | Cysteine3-A | Cysteine3-B | Cysteine3-C |
|---|---|---|---|---|---|---|
| MAG30 (*Vibrio*) | *cyuA* | 28 | 64 | 0 | 2 | 0 |
| MAG30 (*Vibrio*) | malY | 0 | 0 | 0 | 0 | 0 |
| MAG30 (*Vibrio*) | *cysK* | 0 | 0 | 0 | 0 | 0 |
| MAG30 (*Vibrio*) | sseA | 0 | 0 | 0 | 0 | 0 |
| MAG30 (*Vibrio*) | metC | 2 | 0 | 2 | 8 | 2 |
| MAG30 (*Vibrio*) | tnaA | 0 | 0 | 0 | 0 | 0 |
| MAG65 (*Vibrio*) | *cyuA* | 667 | 1195 | 47656 | 117251 | 82272 |
| MAG65 (*Vibrio*) | mgl | 0 | 0 | 28 | 83 | 55 |
| MAG65 (*Vibrio*) | malY | 4 | 9 | 31 | 79 | 62 |
| MAG65 (*Vibrio*) | cysK | 0 | 12 | 220 | 428 | 385 |

| MAG | Gene | Sulfate1-A | Sulfate1-B | Cysteine4 | Cysteine sulfate4 |
|---|---|---|---|---|---|
| MAG249 (*Desulfovibrionaceae)* | *cyuA* | 113 | 15 | 162 | 255 |
| MAG249 (*Desulfovibrionaceae)* | *cysK* | 2 | 0 | 28 | 32 |
| MAG249 (*Desulfovibrionaceae)* | *metC* | 28 | 14 | 0 | 0 |
| MAG249 (*Desulfovibrionaceae)* | *dcyD* | 52 | 2 | 2 | 8 |
| MAG259 (*Desulfovibrionaceae)* | *cyuA* | 2619 | 48 | 37869 | 113461 |
| MAG259 (*Desulfovibrionaceae)* | *cysK* | 541 | 10 | 467 | 803 |
| MAG259 (*Desulfovibrionaceae)* | *metC* | 1338 | 27 | 1226 | 2968 |
| MAG259 (*Desulfovibrionaceae)* | *dcyD* | 117 | 2 | 219 | 312 |
